# Force Gates Regulate Mechanical Transmission Across the Nuclear Envelope

**DOI:** 10.64898/2026.10.06.754136

**Authors:** Nya Domkam, Ghafar Yerima, Shingo Tsukamoto, Mohammad R.K. Mofrad

## Abstract

The LINC (linker of nucleoskeleton and cytoskeleton) complex is a central mechanotransductive apparatus that physically couples cytoskeletal forces to the nuclear interior, thereby regulating nuclear architecture, chromatin organization, and downstream signaling. Using atomistic molecular dynamics simulations, we identify and characterize “force gates” within the LINC complex, structural interactions that maintain mechanical connectivity under load. By introducing targeted mutations into the KASH peptide and SUN protein, we provide computational evidence that a key disulfide bond (SUN2 Cys563 to KASH2 Cys6862) and specific residues within the KASH β strand interaction motif (KASH6875, KASH6877) act as distinct mechanical elements during force transmission across the SUN and KASH interface.

Leveraging this framework, we show that the disulfide bond and the β sheet network contribute unequally: removing the disulfide reduces both sustained and peak force transmission approximately twofold, whereas disrupting the β sheet network leaves peak force largely intact while altering interfacial coupling and the duration over which load is borne. Localized molecular changes therefore reshape mechanical coupling to the nucleus through distinct routes rather than through a single graded loss of force. Such altered LINC mechanics may be relevant to laminopathies, including Hutchinson Gilford Progeria Syndrome (HGPS), in which defective nuclear lamina compromises force transmission; however, implications for specific diseases remain speculative and require experimental validation.

By resolving how distinct molecular contacts contribute to force transmission in space and sequence, this work establishes a framework for dissecting mechanical coupling at protein interfaces under load. These findings provide a foundation for future studies aimed at experimentally testing how individual interface interactions govern nuclear force transmission, a prerequisite for therapeutically targeting LINC mechanics. More broadly, the force gate framework may guide strategies to modulate interfaces that bear mechanical loads in other mechanically regulated proteins. Together, these findings establish a computational framework for investigating force transmission at mechanically loaded protein interfaces and provide experimentally testable predictions for future studies.

## Introduction

Mechanical force is transmitted through cells by defined molecular interfaces, where specific noncovalent and covalent contacts determine how load propagates rather than dissipates^1^. The LINC (linker of nucleoskeleton and cytoskeleton) complex is a paradigmatic example: it spans the nuclear envelope and channels cytoskeletal forces to the nuclear lamina and chromatin through a force-bearing SUN–KASH interface^2,3^. Dysregulation of this mechanical coupling is associated with laminopathies such as Hutchinson–Gilford Progeria Syndrome (HGPS), in which compromised nuclear mechanics drive cellular dysfunction(Fig. 1a)^4,5^.

**Figure 1:**
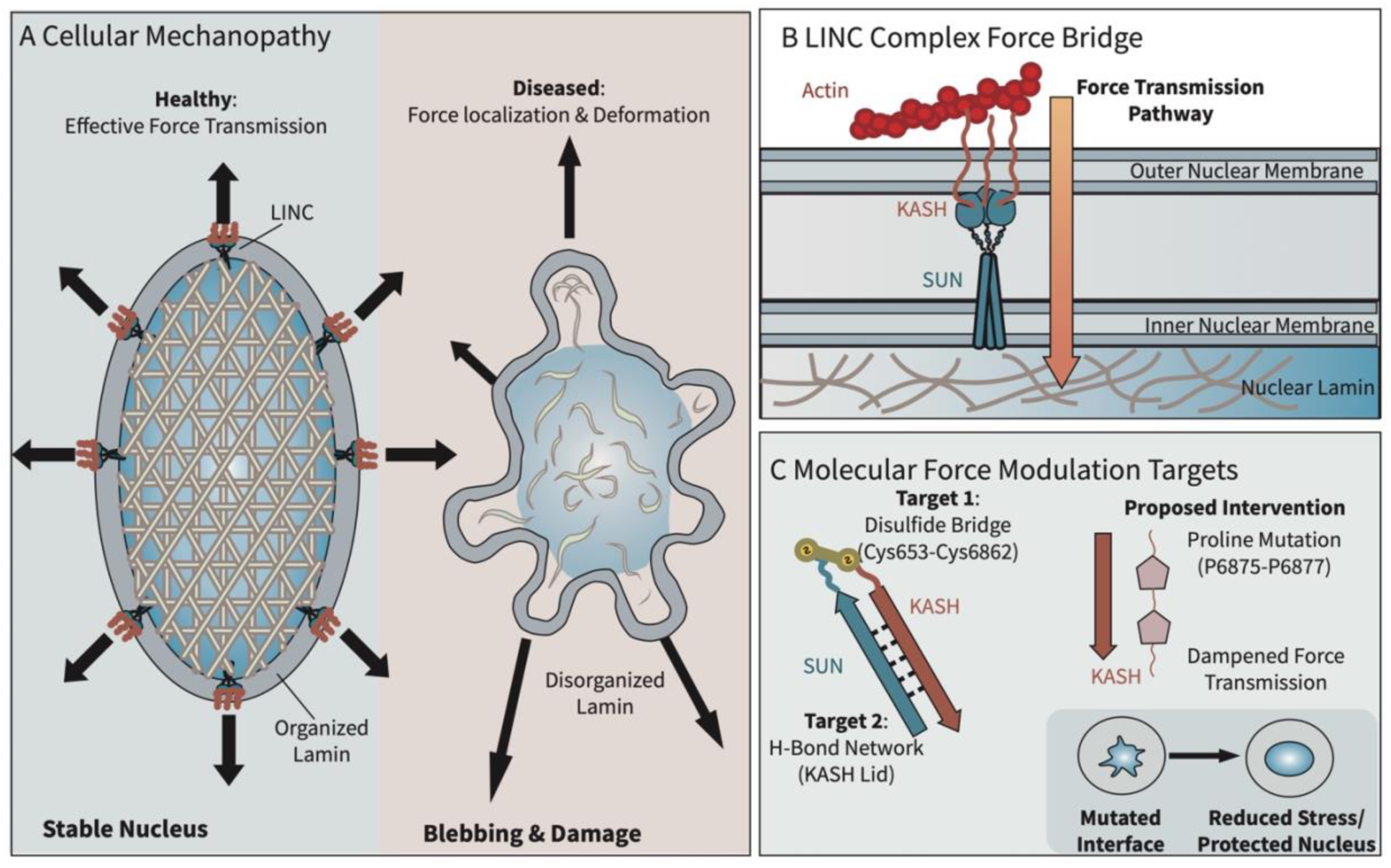
Modulating nuclear mechanics via the LINC complex interface. (A) Schematic comparison of mechanotransduction in healthy and mechanically compromised nuclei (e.g., laminopathies). In healthy cells, forces are distributed across an intact nuclear lamina, whereas in diseased states, compromised lamina structure leads to localized stress, nuclear deformation, and blebbing. *(B)* The LINC complex (SUN and KASH proteins) serves as the primary mechanical linkage between the cytoskeleton and nuclear interior. *(C)* Structural view of the SUN–KASH interface highlighting key interactions investigated in this study, including the disulfide bond and β-sheet hydrogen-bond network. These interactions are hypothesized to function as “force gates” that regulate transmission of mechanical load.

Within the nucleus, the SUN–KASH interface bears the load transmitted from the cytoskeleton, and its mechanical behavior determines how that force is distributed to the lamina and chromatin rather than dissipated at the envelope (Fig. 1b). Force transmission across this interface is bidirectional, supporting both cytoskeleton-to-nucleus signaling and nuclear positioning^6^. Resolving how load propagates through the interface requires atomic-level detail that is difficult to access experimentally; molecular dynamics provides direct access to these interactions under applied force. Here, we use molecular dynamics simulations to identify the structural elements that govern force transmission across the SUN–KASH interface and to determine how their disruption alters mechanical coupling.

The LINC complex consists of SUN (Sad1/UNC-84 homology) proteins in the inner nuclear membrane and KASH (Klarsicht/ANC-1/Syne homology) peptides in the outer nuclear membrane. SUN domains assemble as trimers that can further oligomerize into higher-order assemblies and couple to the lamina through their N-terminal domains^2,3,4^. Mammals express five SUN and six KASH proteins, but only SUN1 and SUN2 are broadly expressed in somatic tissues; the remaining SUN isoforms are testis- restricted^6^. Force transmission through the SUN2–KASH2 pair is mediated by three structural elements at the interface: (1) a disulfide bond between SUN Cys563 and KASH Cys6862, (2) the highly conserved KASH PPPX motif, and (3) β-sheet hydrogen bond network between the KASH peptide and the SUN ’KASH lid’, a SUN loop (residues ∼566–586) that clamps over the bound KASH strand^7,8^.

Disruption of LINC-mediated force transmission is a recurring feature of laminopathies, though the molecular participants differ by disease model and phenotype. In progerin-expressing and *Lmna*-mutant systems, accumulation of SUN1 drives excessive nucleus–microtubule coupling and defective nuclear positioning, and SUN1 depletion rescues these defects^9,10^; in these systems SUN2 levels are reported to be unchanged^9^. In the *Zmpste24⁻/⁻* model under mechanical load, by contrast, SUN2 is elevated and mediates the increased nuclear stiffness, blebbing, and DNA damage characteristic of progeria, with SUN2 suppression restoring nuclear deformability^11^. These observations indicate that both SUN paralogs contribute to laminopathy mechanics through distinct, model-specific mechanisms, with SUN2 specifically implicated in the force-transmission and nuclear-deformation phenotype most relevant to the present study. We therefore examine the SUN2–KASH2 complex, as a tractable atomistic model, chosen for continuity with our group’s prior SUN2–KASH2 force-transmission studies^7,12,13^ rather than because it is the only resolved SUN–KASH structure — SUN1–KASH structures, including higher-order assemblies, have also been reported^2^.

To dissect these contributions, we introduced targeted mutations into the SUN2–KASH2 interface and applied force in molecular dynamics simulations (Fig. 1c). The SUN KASH-lid forms a β-hairpin whose inner strand pairs with the KASH peptide through a backbone hydrogen-bond network^7^. We substituted two KASH β-strand residues, KASH6875 and KASH6877, with proline, which cannot donate a backbone hydrogen bond; these substitutions prevent the KASH strand from registering against the SUN lid, so the inter-strand network fails to form while the overall fold is preserved (table 1). We additionally examined the single substitutions R6877P and M6875P (Fig. S1). Separately, we removed the SUN Cys563–KASH Cys6862 disulfide bond to assess its individual contribution. Prior simulations from our group showed that this disulfide is required for maximal force transmission and that KASH length influences mechanical output^7,12,13^; however, these studies treated the disulfide largely as a binary determinant and did not resolve the intermediate mechanical states through which force propagates. By disrupting the β-sheet network while leaving the disulfide intact, we test whether the disulfide is sufficient for force transmission or whether the β-sheet contributes separably. We did not mutate the conserved C-terminal KASH PPPX motif: this motif inserts into the SUN binding pocket and is required for SUN–KASH complex formation^14^, so its disruption would abolish assembly and preclude analysis of the intermediate mechanical states that are the focus of this work. Together, these perturbations resolve how individual molecular contacts contribute to force transmission across a SUN–KASH interface, establishing a framework for dissecting mechanical coupling at force-bearing protein interfaces more broadly. Throughout this work, we use the term *force-gate framework* to denote the conceptual model of force propagation, and *force gate* as the specific molecular interactions that regulate mechanical load transfer within that framework.

**Table 1.**
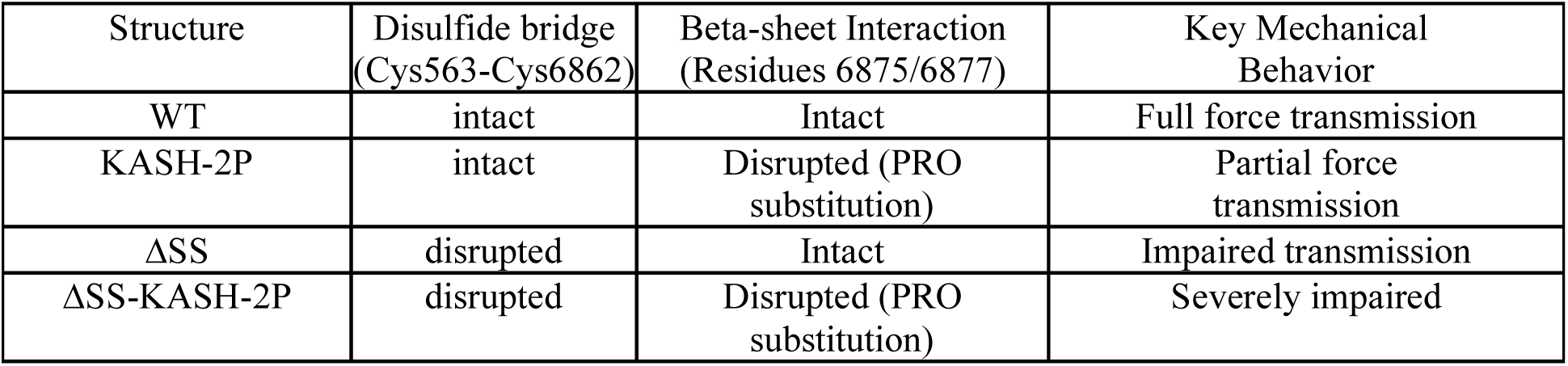
Summary of simulated constructs and associated mechanical features.

## Results

### Point mutations generate structurally stable SUN–KASH assemblies

To evaluate how specific SUN– KASH interactions contribute to LINC mechanics, we introduced point mutations using Discovery Studio^15^ and equilibrated each system for 100 ns of molecular dynamics. Structural stability was assessed by monitoring backbone RMSD and system potential energy (Fig. 2a–c). All systems included a positional restraint on the nitrogen atom on each SUN protomer to preserve biologically relevant geometry consistent with the full-length LINC assembly. Importantly, establishing structural stability across all constructs ensures that subsequent differences in mechanical response arise from specific disruption of force-gating interactions rather than global structural instability.

**Figure 2:**
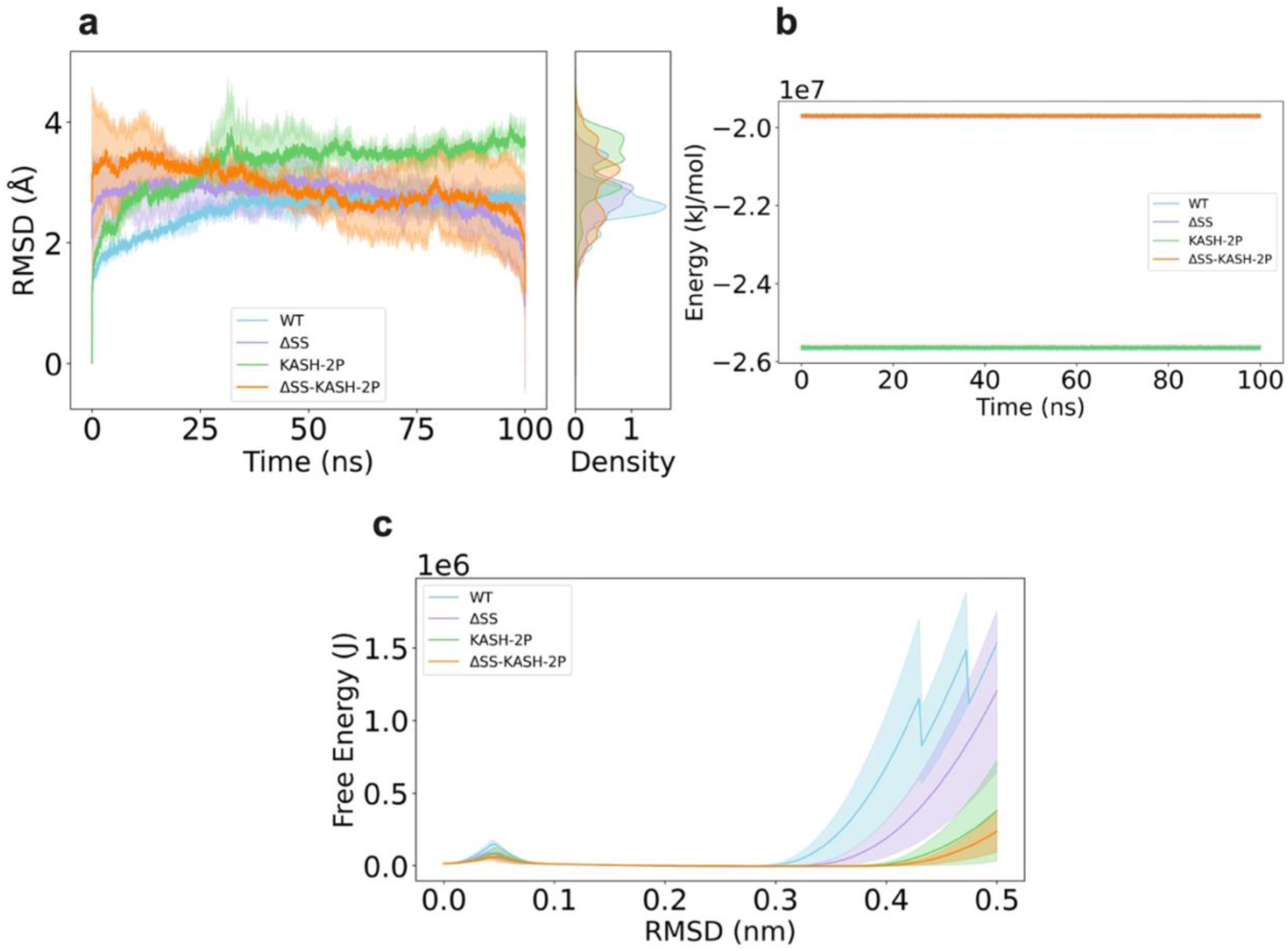
Structural stability of SUN–KASH constructs following equilibration. (a) Backbone RMSD of all constructs over 100 ns of molecular dynamics simulations, with (b) corresponding kernel density distributions. WT exhibits the lowest deviation, followed by ΔSS, while KASH-2P and ΔSS-KASH-2P show higher structural variability due to proline-induced disruption. (c) Total system potential energy (protein, solvent, ions), showing lower energies for constructs retaining the disulfide bond. (d) One-dimensional free energy landscapes as a function of RMSD. WT displays the deepest minimum, indicating highest stability. These results confirm that all constructs reach stable conformational ensembles suitable for mechanical analysis.

As expected, the wild-type (WT) structure exhibited the lowest RMSD and most favorable potential energy, reflecting its native stability. Removal of the SUN2 Cys563–KASH2 Cys6862 disulfide bond (ΔSS) resulted in moderately increased RMSD, consistent with enhanced conformational freedom of the KASH peptide once uncoupled from SUN. Introduction of the two proline substitutions (KASH-2P and ΔSS- KASH-2P) further elevated RMSD values, in agreement with the known secondary-structure-disrupting effect of proline^16^. The slightly higher RMSD observed in KASH-2P relative to ΔSS-KASH-2P may arise from backbone strain imposed by maintaining the disulfide constraint alongside the disruptive prolines. Importantly, RMSD in all systems plateaued within ∼10 ns, indicating that each construct reaches a stable conformational ensemble.

Potential energy analysis provided a complementary measure of stability (Fig. 2c). Systems retaining the disulfide bond consistently displayed lower (more favorable) total potential energy than their ΔSS counterparts, though the magnitude of the difference likely reflects both covalent linkage and solvation/conformational rearrangements beyond the bond itself. To further resolve intrinsic protein stability, we computed 1D free-energy profiles as a function of RMSD (Fig. 2d). WT again showed the deepest minimum, followed by ΔSS, ΔSS-KASH-2P, and KASH-2P—an ordering that differs from total potential energies due to the latter including solvent contributions.

Together, these analyses confirm that all engineered variants adopt **stable structural states**, enabling direct comparison of their mechanical behavior in subsequent force-loading simulations.

### Proline mutations disrupt β-sheet interactions and secondary-structure integrity at the SUN–KASH interface

To determine how proline substitutions alter the SUN–KASH interface, we analyzed φ/ψ backbone dihedral angles and hydrogen bonding within KASH residues 6874–6879 (Fig. 3). Wild-type (WT) and ΔSS structures displayed nearly identical φ/ψ distributions (Fig. 3a,b), exhibiting predominantly β-strand character with occasional right-handed α-helix sampling. Consistent with preserved KASH-lid engagement, both constructs spent >91% of simulation time in favorable Ramachandran regions (WT: 91.35%; ΔSS: 92.23%). Slightly increased conformational sampling in ΔSS likely reflects the increased mobility of KASH when not covalently tethered to SUN via the disulfide bond.

**Figure 3:**
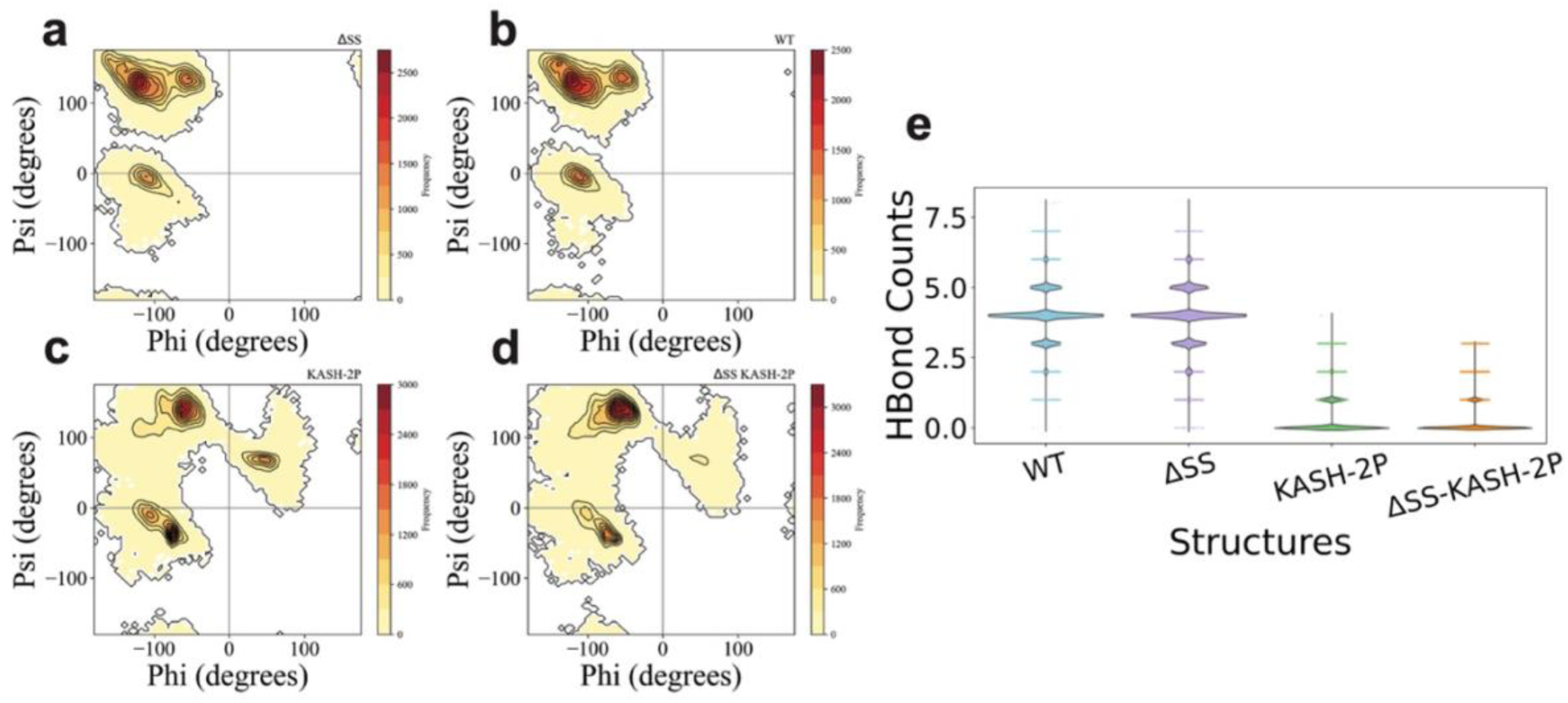
Disruption of β-sheet structure by proline mutations. Ramachandran plots for (A) ΔSS, (B) WT, (C) KASH-2P, and (D) ΔSS-KASH-2P. WT and ΔSS retain β-strand conformations, while proline substitutions shift distributions toward polyproline- like regions and increase sampling of unfavorable conformations. (E) Hydrogen-bond analysis between KASH residues (6870–6878) and the SUN KASH-lid (578– 586) shows that WT and ΔSS maintain stable β-sheet interactions, whereas proline mutants exhibit near-complete loss of hydrogen bonding. These results confirm disruption of the β-sheet force-gate at the SUN–KASH interface.

In contrast, the double-proline mutations substantially altered backbone geometry (Fig. 3c,d). Both KASH-2P and ΔSS-KASH-2P showed marked decreases in favorable conformations (75.46% and 79.09%, respectively) and increased sampling of acceptable/unfavorable regions (including 13.05–15.73% in unfavorable space). These shifts are consistent with the steric constraints of proline and its inability to donate backbone hydrogen bonds^17^. The φ/ψ distributions exhibited clear enrichment near −75° φ / −50° ψ, characteristic of polyproline-like conformations^17^, and reduced occupancy of the β-sheet quadrant. Notably, ΔSS-KASH-2P exhibited slightly broader conformational sampling than KASH-2P, consistent with increased flexibility when the disulfide constraint is removed.

Hydrogen-bond analysis confirmed this structural disruption: WT and ΔSS retained persistent KASH-lid hydrogen bonding, whereas double-proline mutants showed a near-complete loss of SUN– KASH β-sheet interactions (Fig. 3e). This validates the intended functional effect of the mutation physically uncoupling the KASH peptide from SUN’s lid region.

Together, these results support that proline substitutions effectively break the KASH β-strand force-bearing interface, reducing secondary-structure integrity and altering local conformational dynamics in a manner expected to impair mechanical load transfer.

### Energetic decomposition reveals loss of SUN–KASH coupling in proline mutants

To quantify how mutations alter the binding energetics between SUN and KASH, we performed per-residue free-energy decomposition using gmx_MMPBSA^18,19^. This method estimates effective interaction energy as the sum of molecular mechanics gas-phase terms and solvation contributions:

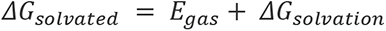

Analysis focused on the β-strand interaction interface spanning KASH6870–6878 and SUN573– 586 (Fig. 4,5). WT and ΔSS structures showed similar energetic profiles (Fig. 4a,b), consistent with preserved hydrogen-bonding interactions and comparable secondary structure in both cases. In contrast, double-proline substitutions led to substantial reductions in stabilizing interaction energy at SUN LEU573 and SUN SER575, residues that normally form hydrogen bonds with KASH6875 and KASH6877. Because proline lacks a backbone amide hydrogen, the β-sheet interface cannot form in either KASH-2P or ΔSS- KASH-2P, causing the KASH strand to disengage from the lid.

**Figure 4:**
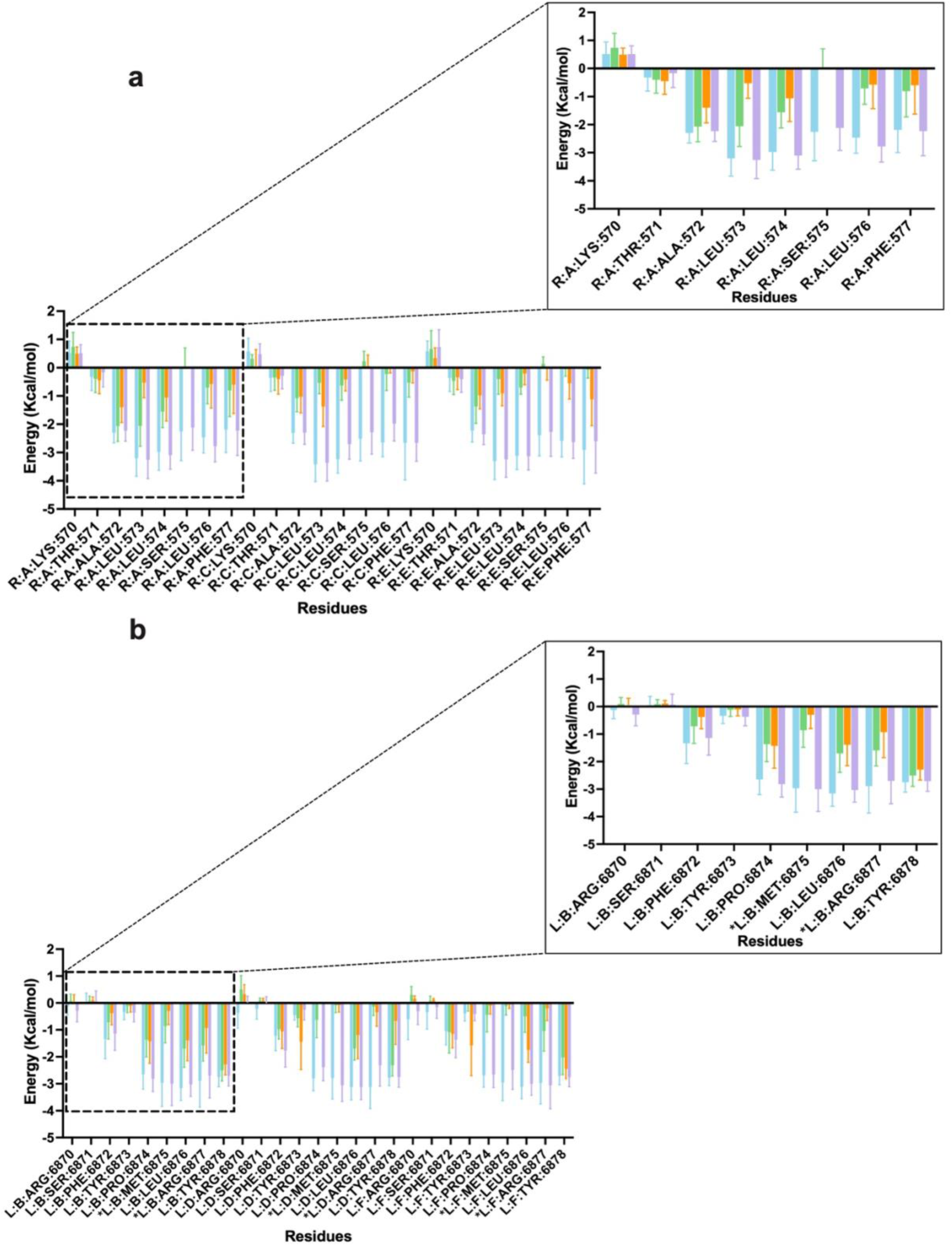
Energetic consequences of β-sheet disruption. (A) Per-residue energy decomposition of the SUN KASH-lid region shows reduced stabilizing interactions in proline mutants. (B) Energy decomposition of the KASH peptide highlights altered energetics at mutation sites (6875, 6877).

**Figure 5:**
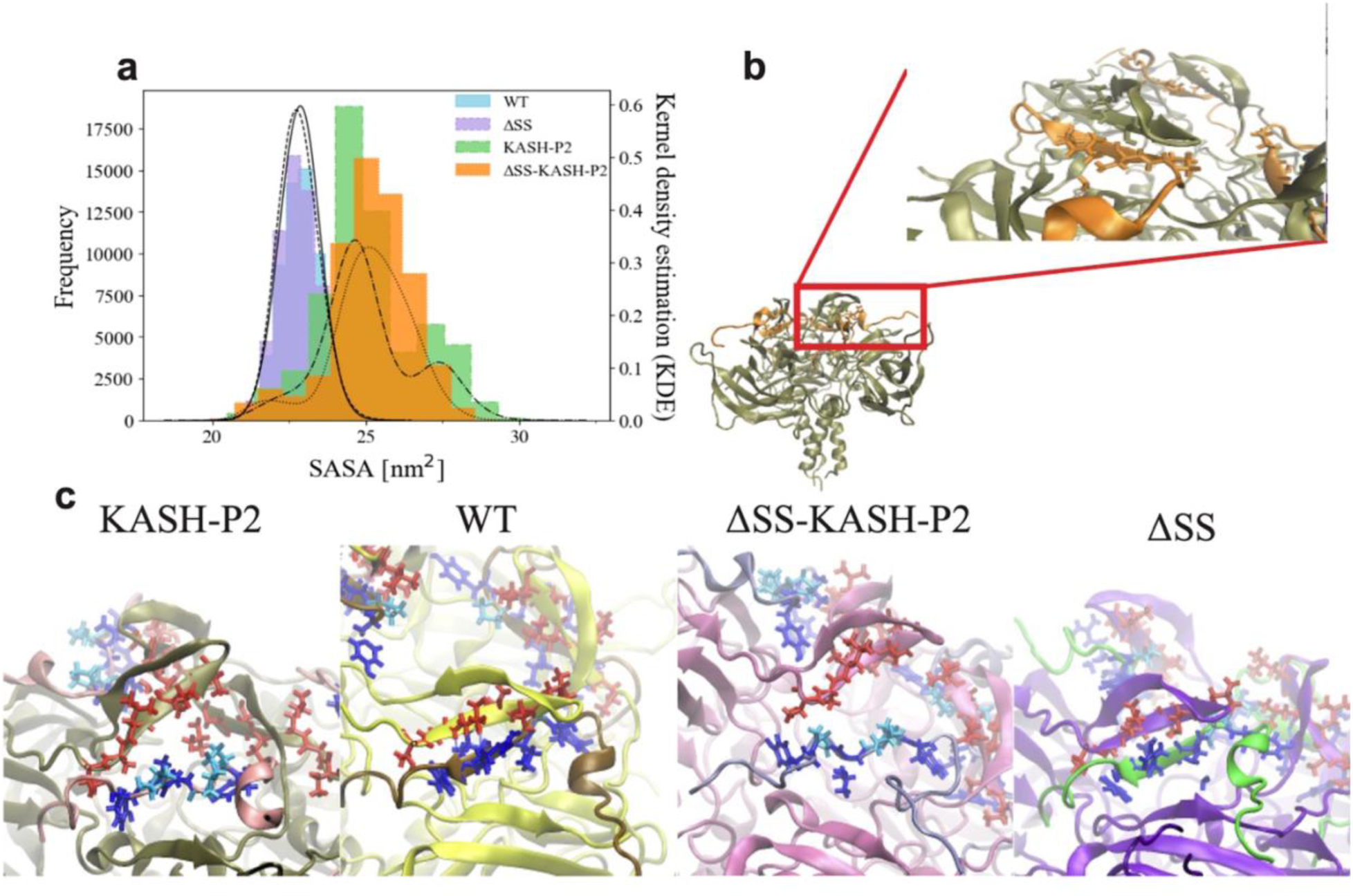
Structural consequences of β-sheet disruption. (A) Solvent-accessible surface area (SASA) of KASH increases in proline mutants, indicating loss of burial at the interface. (B–D) Structural snapshots show separation between KASH and the SUN lid in mutant constructs. Together, these results support weakening of SUN–KASH coupling upon disruption of the β- sheet force gate.

These energetic trends align with solvent-exposure measurements (Fig. 5a). WT and ΔSS exhibit nearly identical SASA values, whereas both mutant structures show significantly increased solvent accessibility, indicating a loss of SUN-KASH burial and packing at the interface. Slightly lower SASA in KASH-2P relative to ΔSS-KASH-2P reflects the disulfide bond’s continued geometric constraint preventing larger-scale separation.

Interestingly, several tyrosine residues in the KASH peptide of ΔSS-KASH-2P display unusually elevated interaction energies relative to all other systems. We attribute this effect to cation–π and π–π contacts that transiently stabilize partially detached KASH chains, particularly when the absence of a disulfide constraint allows deeper sampling of alternative conformations. These interactions likely represent transient compensatory contacts that are mechanically weaker than the native β-sheet interface. Collectively, these results support the interpretation that β-sheet interactions act as a critical force gate governing interfacial stability.

Collectively, the energetic landscape confirms that proline substitutions disrupt native binding energetics, weakening SUN–KASH coupling and increasing solvent exposure in a manner directly consistent with loss of a continuous force-transmission pathway.

### Cross-correlation analysis identifies loss of coordinated force propagation in mutant constructs

To evaluate how mutations alter force propagation through the LINC complex, we performed residue–residue cross-correlation analysis during both equilibrium (production) and force-loaded (steered pulling) simulations (Fig. 6a,b). This approach identifies regions in which residues move cooperatively under mechanical load, reflecting their participation in a continuous force-bearing network.

**Figure 6:**
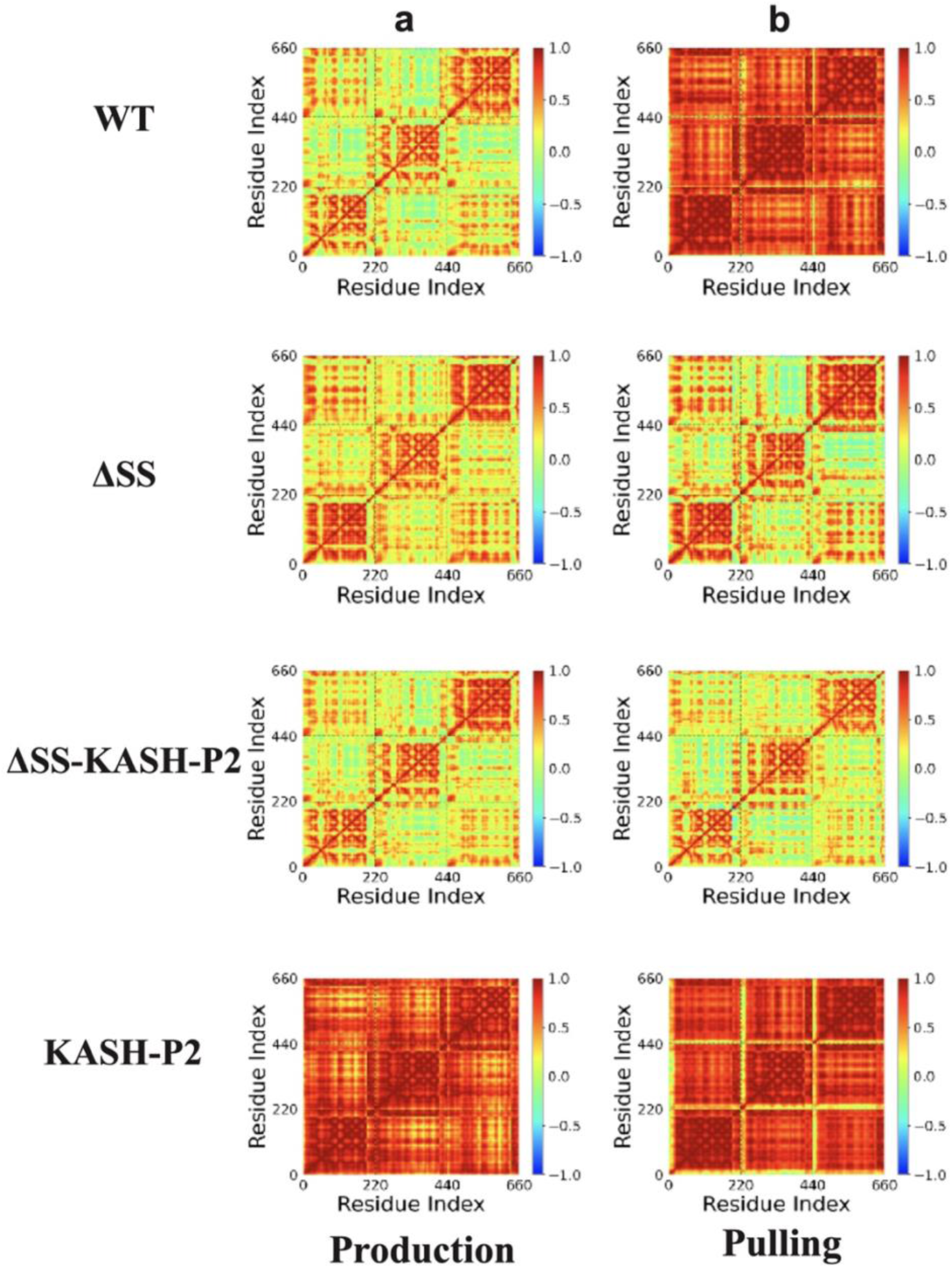
Mechanical response of LINC constructs. (A–B) Residue cross-correlation matrices during equilibrium (A) and pulling (B). WT exhibits strong long-range correlations, while ΔSS constructs show reduced inter-protomer coupling.

Wild-type structures exhibited strong positive correlations across protomers during pulling simulations, indicating coordinated motion throughout the SUN trimer. In contrast, mutants lacking the SUN–KASH disulfide bond (ΔSS and ΔSS-KASH-2P) showed extensive regions with little or no correlation, particularly between different protomers. These results suggest that loss of the disulfide tether prevents forces from propagating beyond the immediate SUN–KASH interaction site, isolating domains that would normally deform together.

Structures retaining the disulfide bond (WT and KASH-2P) maintained higher levels of long- range coordination, although KASH-2P displayed reduced coupling near the KASH lid, consistent with β- sheet disruption. Notably, KASH-2P’s cross-correlation heatmap during production simulations already resembled the pulling response of ΔSS constructs, indicating that proline mutations alone are sufficient to weaken baseline mechanical connectivity.

To more precisely identify mechanically responsive regions, we compared cross-correlation differences between production and pulling conditions. In the WT complex, multiple SUN regions exhibited strong correlation increases under load, consistent with engagement of multiple force-transmission pathways. By contrast, mutant structures showed strong correlation changes localized almost exclusively to the KASH peptide, reflecting restricted force transfer and early mechanical isolation.

Together, these data demonstrate that both the β-sheet interface and the disulfide bridge are essential for global motion coupling, and that their disruption yields measurable “break points” in mechanical communication within the LINC complex. These altered correlation signatures directly predict changes in mechanical performance, which we next evaluate using force-loading simulations and point- change detection analysis.

### Force-gate identity determines the magnitude and duration of force transmission

To directly assess how disruption of specific force gates affects mechanical performance, we analyzed force–time trajectories alongside structural transitions (Fig. 7a-d). Across all six constructs, both trajectory-mean and peak force separated disulfide-retaining from disulfide-deleted systems without overlap of replicate standard deviations (mean: 348–446 pN vs 142–190 pN; peak: 626–756 pN vs 381–460 pN; Fig. S1c,d). No comparison within either group produced a separation of comparable size.

**Figure 7:**
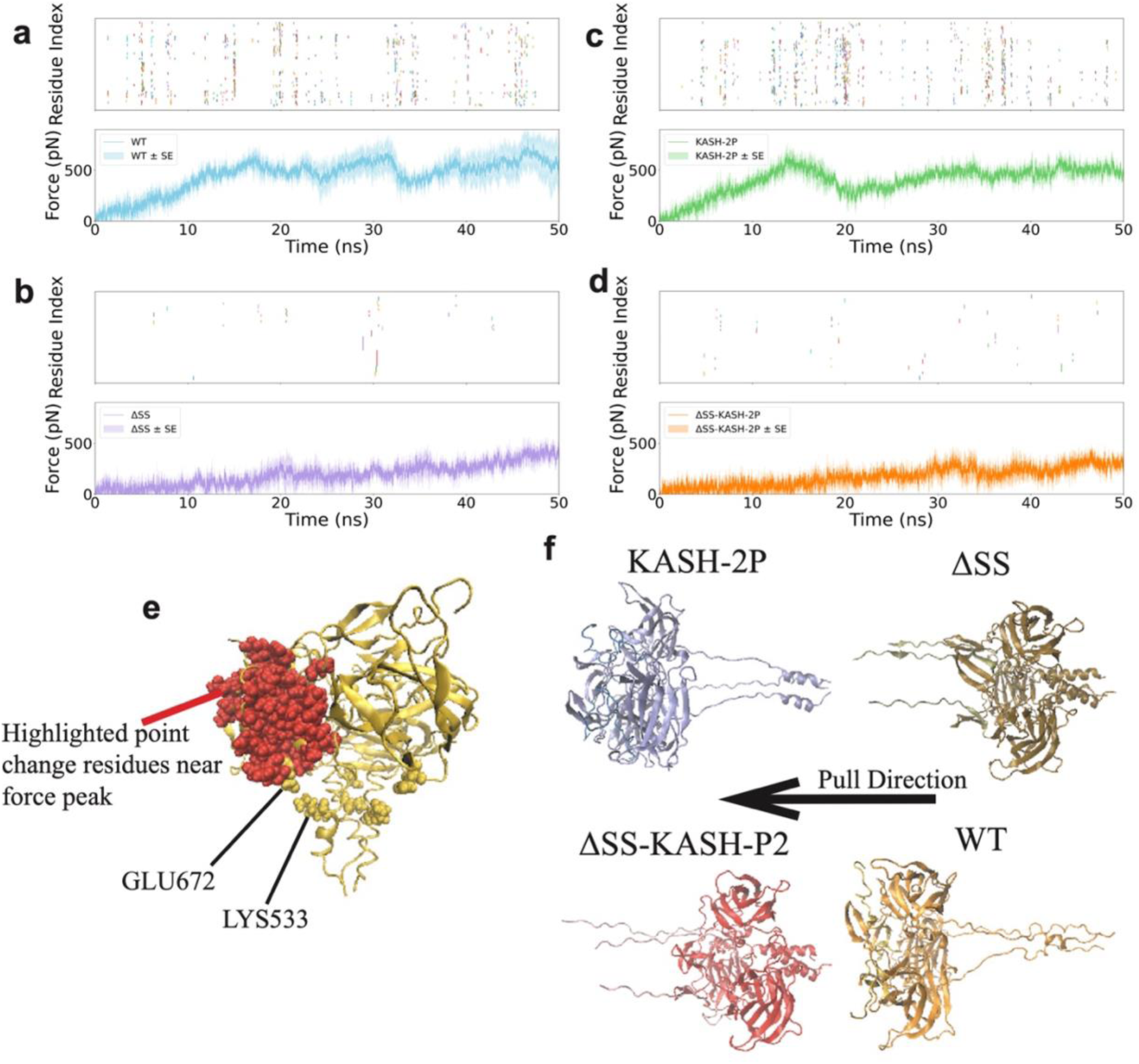
Force propagation of LINC constructs. (A–D) Point-change detection maps aligned with force–time traces for WT, ΔSS, KASH-2P, and ΔSS-KASH-2P. WT shows the highest density of coordinated transitions, while mutants display progressively reduced coupling. (E) Structural mapping of residues undergoing coordinated transitions at peak force. (F) Final conformations under load highlight distinct deformation modes depending on force- gate integrity. These results suggest that disruption of specific interactions alters global force transmission pathways.

The WT complex, which retains the disulfide bond and the β-sheet hydrogen-bond network, and, by inference from prior work, the downstream salt-bridge contacts ^7^, sustained the highest forces and exhibited broad force distribution across the trimeric assembly. KASH-2P, which retains the disulfide bond but lacks the β-sheet gate, showed lower mean and peak force than WT (mean 396 ± 13 vs 446 ± 76 pN; peak 626 ± 70 vs 756 ± 200 pN), though in both metrics the difference falls within the replicate spread of WT and is not resolved at n = 3. The force–time traces are more informative here: KASH-2P tracks WT through the early loading phase and diverges later in the pull (Fig. 7a,c; S1a,b), indicating that loss of the β-sheet network alters where in the trajectory force is sustained rather than eliminating the interface’s capacity to bear load. Both ΔSS and ΔSS-KASH-2P, which lack the disulfide bond, showed substantially lower maximum forces, consistent with the disulfide gate being required for force propagation to reach beyond the immediate SUN–KASH interface. ΔSS-KASH-2P showed lower mean and peak force than ΔSS (mean 170 ± 37 vs 190 ± 14 pN; peak 381 ± 58 vs 460 ± 36 pN), but the reduction is small relative to the effect of removing the disulfide and, in the trajectory mean, falls within replicate spread. Once the disulfide is removed, additional disruption of the β-sheet network produces at most a modest further loss.

The single-proline constructs dissociate peak capacity from sustained transmission. In each background, the single and double substitutions reached comparable peak forces (M6875P 666 ± 68 vs KASH-2P 626 ± 70 pN; R6877P 382 ± 38 vs ΔSS-KASH-2P 381 ± 58 pN) while differing in trajectory mean (348 ± 13 vs 396 ± 13 pN; 142 ± 7 vs 170 ± 37 pN). The force–time traces locate this difference in the loading phase: M6875P reaches high force later than KASH-2P (Fig. S1a,b) and then sustains comparable force. A single β-strand substitution therefore shortens the interval over which the interface bears load without lowering the maximum load it can sustain.

Point-change detection analysis reveals discrete, reproducible transitions in the force–time profiles that correspond to specific structural rearrangements at the SUN–KASH interface. In this approach, inspired by Iakhiaev 2013^20^, residue positions throughout the pulling trajectory are treated as time-series signals, and kernel-based change-point detection identifies moments at which residues undergo significant coordinate shifts. Residues exhibiting simultaneous point changes are interpreted as being mechanically connected through a shared force-transmission pathway. The WT structure displayed the highest density of point changes across both time and residue space, consistent with forces traversing multiple gates and engaging a broad network of residues. Fewer point changes were observed in constructs lacking one or more force gates, with ΔSS-KASH-2P showing the fewest. KASH-2P was a partial exception to this trend: its cross-correlation heatmap during production simulations closely resembled its pulling response, suggesting that proline-induced disruption of the β-sheet gate alters baseline mechanical connectivity even before external load is applied.

Critically, point-change detection maps can be directly overlaid onto force-versus-time curves, enabling residue-level mapping of structural transitions to specific force events. This correspondence reveals that each passage through a force gate produces a characteristic signature in the force trace, providing a mechanistic link between molecular interactions and macroscopic force output. These mappings were visualized as annotated trajectories in VMD^21^.

Together, these results indicate that force transmission through the LINC complex is mediated by discrete molecular contacts whose disruption reduces the magnitude of transmitted forces, consistent with the gate-like engagement.

## Discussion

The LINC complex transmits cytoskeletal force across the nuclear envelope, shaping nuclear architecture, chromatin organization, and downstream signaling^22˒23^. Disruption of LINC mechanics is implicated in laminopathies such as HGPS, where abnormal force transmission contributes to nuclear deformation and genomic instability. Our simulations demonstrate that the β-sheet network and disulfide bond contribute separably and through different observables. The disulfide sets the magnitude of transmitted force: its removal halves both sustained and peak force. The β-sheet network governs interfacial coupling and the duration of engagement: its disruption is evident in hydrogen bonding, solvent exposure, per-residue energetics, and cross-correlation, yet leaves peak force within replicate spread of WT.

Although we did not systematically analyze all single-point mutations in the main text, the behavior of partially disrupted constructs is consistent with a response that is bimodal with respect to the disulfide and graded within each disulfide state. This observation supports the idea that SUN–KASH force transmission depends on multiple structurally distinct interactions that contribute unequally to mechanical stability^8^.

Building on prior work showing that KASH-peptide variation alters LINC force responses^7,13^, we propose a force-gate framework in which discrete molecular interactions function as force gates that regulate the propagation of mechanical load (Fig. 8a). Throughout this work, we use the term force-gate framework to denote the conceptual model of force propagation and force gates to denote the specific molecular interactions that regulate mechanical load transfer. Rather than simply cataloguing mechanically important residues, this framework distinguishes force gates according to when and how they engage during force propagation. Our simulations support a model in which force transmission through the SUN–KASH interface proceeds through distinct, ordered events rather than a single all-or-none rupture: partial decoupling, in which some interactions fail while reduced force transmission persists, and more complete decoupling, in which loss of a primary contact prevents force from reaching downstream elements. By relating point-change-detection maps to force–time profiles, we identify which interactions contribute to force propagation and the approximate order in which they engage.

**Figure 8.**
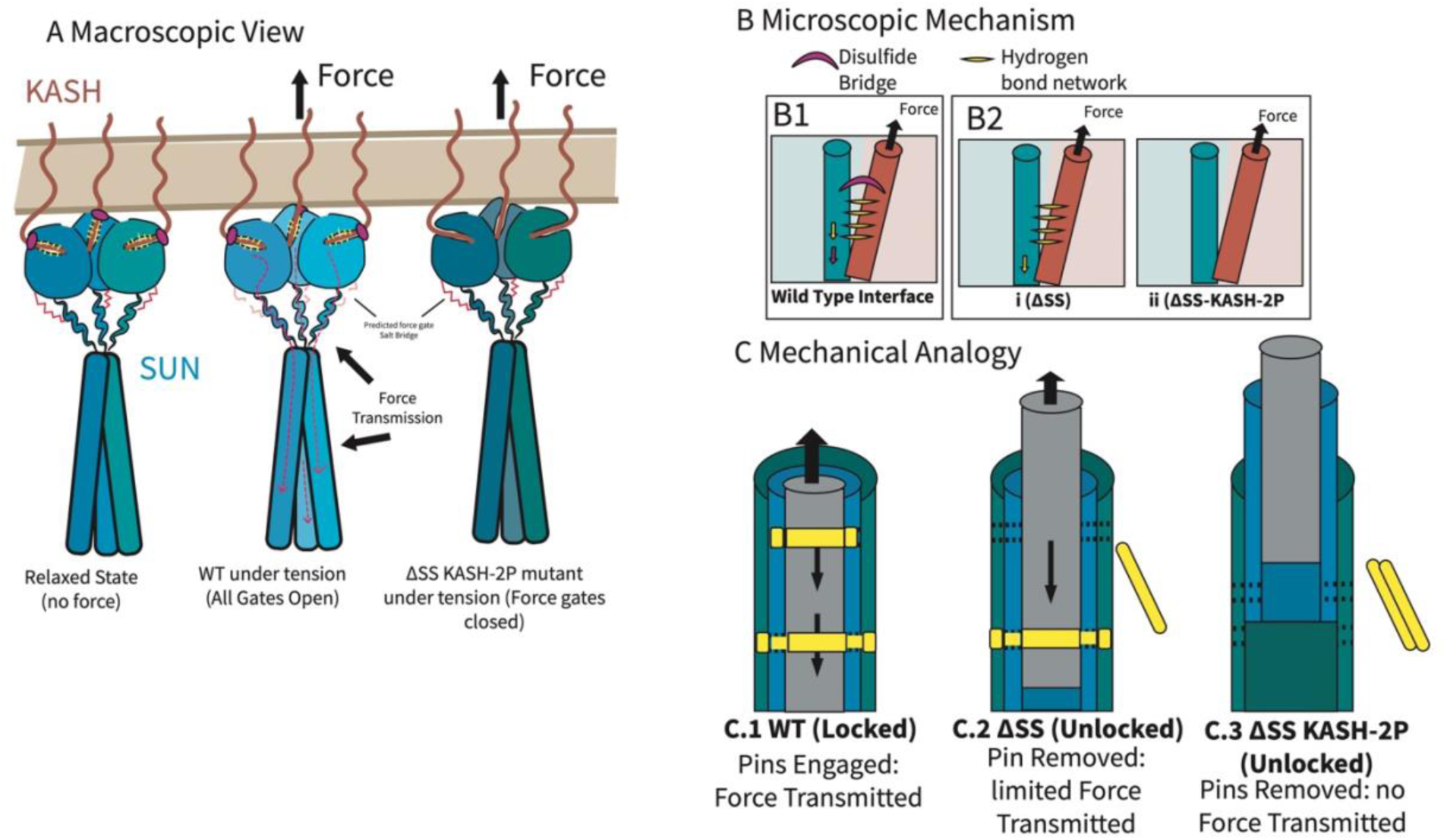
Schematic summary of the force-gate framework in the LINC complex. (A) Conceptual view of force transmission through the SUN–KASH complex. In the WT complex, applied tensile force propagates through the interface and into the SUN trimer. In mutants lacking one or more force gates, force transmission is attenuated or arrested at the SUN–KASH boundary. *(B)* Molecular view of the primary force-gating interactions. The disulfide bridge and β-sheet hydrogen-bond network cooperate to transmit force across the SUN–KASH interface. Disruption of either interaction alters the force-propagation pathway. *(C)* Mechanical analogy illustrating sequential force gates. The LINC complex behaves as a mechanically coupled system in which disruption of individual gates progressively reduces force transmission efficiency.

Within the SUN–KASH system, our simulations provide direct computational evidence that two molecular interactions function as force gates within the proposed framework: the disulfide bond between SUN2 Cys563 and KASH2 Cys6862, and the β-sheet hydrogen-bond network between the SUN KASH-lid and the KASH peptide (Fig. 8b). When both are intact, force is distributed across the trimeric assembly and the interface withstands higher loads; disrupting the β-sheet network or removing the disulfide reduces maximum force resistance and produces more localized structural responses. These complementary analyses consistently support this interpretation. Based on prior work, we further propose a third downstream candidate force gate involving salt-bridge coupling in the coiled-coil/β-barrel region (Fig. 8c). We did not perturb this interaction here, but prior simulations from our group showed that these salt bridges rupture under load^7^, supporting the hypothesis that they represent an additional force gate requiring direct experimental or computational testing.

The force-gate framework may extend to other LINC complexes, although the specific force gates are likely to differ. The force-gate framework may be conserved at the level of interaction *type* — covalent, hydrogen-bonding, or electrostatic — while varying in molecular detail between systems: the disulfide characterized here is not universal across SUN–KASH pairs, and salt-bridge organization differs across SUN isoforms, whereas the KASH-lid architecture appears more broadly conserved. The identity, strength, and ordering of individual gates must therefore be determined per system. In living cells, SUN2 and nesprin- 2 approach a 3:3 composition only as a limiting state: dual-color tsMSQ in the nuclear envelope gives brightness values rising from 1 toward 3 with expression level, reaching the trimeric limit only in cells with nesprin-2:SUN2 ratios above 4:1^32^. Full occupancy of the three KASH sites is therefore attainable but not typical. Under such substoichiometric occupancy — partial occupancy of the three KASH sites on a SUN2 trimer — gate engagement is expected to be heterogeneous, with only load-bearing pairs engaging while unoccupied or unloaded sites remain mechanically inactive, a prediction this study does not test.

The SUN–KASH interface occupies a distinct position in the cellular force-transmission pathway. Most current approaches to mechanotransduction act at the cell–matrix interface, through integrins, talin, or actomyosin signaling, whereas the contacts examined here operate at the nuclear envelope itself, where cytoskeletal load is relayed to the nuclear interior.

Several limitations of this study should be noted. First, the conclusions are derived from atomistic simulations and therefore require experimental validation. Second, pulling rates in steered molecular dynamics exceed those typically used in single-molecule experiments, so absolute rupture forces should be interpreted cautiously. Since peak force is a single-timepoint estimate, it carries greater replicate variance than the trajectory mean and should be compared across constructs with correspondingly more caution. Although the relative sequence of force-gate failure appears qualitatively consistent across replicates, quantitative force thresholds may depend on loading rate. Third, while our simulations directly support force-gating roles for the disulfide bond and the β-sheet hydrogen-bond network, the contribution of salt bridges remains a model-based inference rather than a directly perturbed variable in this study. Fourth, force was applied symmetrically to all three KASH peptides. If the three sites are only partially occupied in cells, loading would be asymmetric and distributed over fewer engaged pairs; the simulations therefore describe the fully loaded reference case rather than the typical in-cell configuration. Finally, the applicability of this framework to other LINC isoforms remains to be tested explicitly.

Several directions follow from this work. The force-gate framework yields concrete, testable predictions: that the disulfide and the β-sheet network are distinguishable by the observable they affect, force magnitude versus engagement duration, and that partial disruption within a single gate reduces the interval of load-bearing without lowering peak capacity. These differences are reflected in the differential mechanical stability of LINC mutants under load. Since point-change analysis provides per-residue mechanical sensitivity, similar analyses could in principle be applied to identify force-bearing contacts in other mechanically loaded protein interfaces. Whether interface contacts of the kind examined here could ultimately be modulated to tune nuclear force transmission for therapeutic benefit remains a speculative and untested possibility, contingent on experimental validation and on extending these analyses beyond the single, truncated SUN–KASH pair studied here.

In conclusion, by proposing and evaluating a force-gate framework for the LINC complex, we provide a mechanistic basis for how discrete molecular interactions regulate nuclear force transmission and establish a foundation for future experimental studies.

## Methods

Computational models of the SUN2–KASH2 trimer were constructed from the crystal structure with PDB ID 4DXS^24^. We did not model the full coiled-coil luminal domain of SUN2: AlphaFold2 and AlphaFold3 predict the extended α-helical coiled-coil inconsistently, and the associated per-residue confidence scores in this region are too low to provide a reliable basis for force simulation. Higher-order SUN–KASH assemblies and the full luminal domain may redistribute load differently than the truncated construct modeled here^2^, a limitation we return to in the Discussion. Mutant constructs were generated in BIOVIA Discovery Studio^15^ by substituting KASH residues 6875 and 6877 with proline, to disrupt β-strand hydrogen-bonding interactions at the KASH lid while maintaining peptide continuity. The disulfide- removed construct (ΔSS) was generated by removing the disulfide-bond connectivity between Cys563 and Cys6862 while retaining both cysteine residues; no residue substitutions were made. Here, ΔSS denotes loss of the disulfide bond, not a serine substitution. The structural models were subsequently prepared for molecular dynamics simulations in GROMACS version 2021^25^.

All simulations employed the CHARMM36 force field^26^ and explicit TIP3P water molecules within a periodic orthorhombic box. Charge neutralization and solvation conditions were designed to approximate the nucleoplasmic environment by adding ions to a final concentration of 0.15 M KCl and 0.05 M CaCl₂^27,28^. Long-range electrostatic interactions were computed using the Particle Mesh Ewald method with a real-space cutoff of 1.2 nm. To reflect the physical continuity of the coiled-coil domain beyond the crystallized segment, a positional restraint was applied to the nitrogen atom of Gly522 of each of the three SUN protomers throughout all simulations.

Energy minimization of each system was conducted using steepest descent for 5,000 steps with a convergence tolerance of 1,000 kJ/mol·nm. Systems were equilibrated in the NPT ensemble for 5 ns using a 1 fs timestep, constant temperature (310 K) and pressure (1 bar) controlled via Berendsen thermostat and barostat, respectively, with heavy-atom positional restraints applied to the protein. Production simulations were then run for 100 ns, during which restraints were limited to the anchored coiled-coil residue, and temperature and pressure were controlled with the Nose–Hoover thermostat and Parrinello–Rahman barostat. Three replicate simulations with randomized initial velocities were performed for each of the six structural variants to confirm reproducibility.

Following relaxation, mechanically induced deformation was assessed using steered molecular dynamics. A constant-velocity pulling protocol was applied to the Oxygen atom of Cys6862 of all three KASH peptide at 1 Å/ns for 50 ns while maintaining the restraint of Gly522 nitrogen on all three SUN protomers as the reference position. Pulling all three KASH peptides against the symmetric SUN anchor reproduces the symmetric loading geometry of the trimeric assembly under tension. This is a deliberate idealization: in cells the three KASH sites may be only partially occupied^32^, in which case loading would be asymmetric and distributed over fewer engaged pairs. Our simulations therefore characterize the fully-loaded reference case rather than any specific substoichiometric configuration. All parameters including solvent, force field, temperature, pressure, and electrostatics were identical to the production simulations.

Force–extension and force–time profiles were extracted from the SMD trajectories by retrieving the pulling force at each timestep from GROMACS output and averaged across the three replicates, with shaded bands denoting the standard error of the mean. Trajectory-mean force was computed as the average pulling force over the full 50 ns of each replicate; peak force was taken as the maximum 1-ns-averaged force within each replicate. Shaded bands on the force–time traces in Fig. 7 denote the standard error of the mean across replicates; shaded bands in Fig. S1a,b and all bar charts show the standard deviation, so that replicate spread rather than precision of the mean is displayed for the summary comparisons.

### Post-Processing/Trajectory analyses

All post-processing and analysis of trajectory data were performed using GROMACS utilities^25^ and custom scripts written in Python^29^. Backbone structural stability was characterized by root mean square deviation (RMSD). Frames from each trajectory were first aligned to the initial minimized structure, and the RMSD of backbone heavy atoms was computed over time. System potential energy was extracted from GROMACS energy files over the production trajectories as an additional stability metric.

Molecular visualizations, including structural snapshots and trajectory renderings, were generated using Visual Molecular Dynamics (VMD)^21^.

Per-residue free-energy contributions were quantified using the gmx_MMPBSA package^18,19^ employing the generalized Born implicit solvent model. Only residue pairs within 6 Å of one another at any point during the simulation were considered for free-energy evaluation to specifically capture the energetic contributions at the SUN–KASH interface. Entropic contributions were omitted based on the large computational cost and their minimal effect on relative binding trends for closely related complexes.

Residue–residue dynamic coupling was quantified using displacement-based cross-correlation analysis. For each residue *i*, the displacement vector relative to the time-averaged position was defined as:

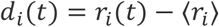

where **r***_i_*(*t*) is the instantaneous position of the Cα atom and ⟨**r***_i_*⟩ is its mean over the trajectory. The normalized zero-lag correlation between residues *i* and *j* was then calculated as:

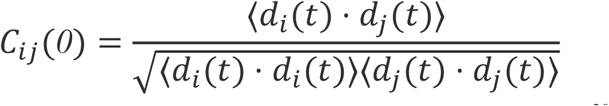

The code base used for this analysis was adapted from the MD-TASK framework^30^ to accommodate the trimeric SUN–KASH assembly.

To identify transitions in mechanical communication under load, residue trajectories were treated as spatial time-series signals for kernel-based change-point detection. Residue positions were projected into a reproducing-kernel Hilbert space using a Gaussian radial basis function kernel. Changes in the mean of the transformed signal were detected to locate molecular force-gate engagement events. These analyses were conducted using the ruptures Python package^31^.

Local conformational effects of mutations were assessed by computing Ramachandran backbone dihedral distributions using the GROMACS “rama” function, with φ and ψ angles plotted in Python. Solvent-accessible surface area (SASA) for the KASH peptide was calculated using the GROMACS “sasa” tool. Hydrogen bonding between the KASH peptide and the SUN KASH-lid interface was quantified by counting donor–acceptor pairs within 0.35 nm and with appropriate angular geometry criterion.

The one-dimensional free-energy landscape was derived from RMSD probability distributions according to the Boltzmann relation:

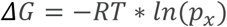

where Δ*G* is the free-energy difference associated with an RMSD state, *R* is the universal gas constant, *T* is the simulation temperature (310 K), and *p_x_* is the normalized probability of observing RMSD value *x* over the trajectory.

## Acknowledgements

This research used the Expanse at San Diego Supercomputer Center through allocation BIO230228 from the Advanced Cyberinfrastructure Coordination Ecosystem: Services & Support (ACCESS) supercomputing facilities, supported by the National Science Foundation (NSF) (grant nos. 2138259, 2138286, 2138307, 2137603, and 2138296). This research was also supported through the QUAD Fellowship. Fruitful discussions with Mohammad Khavani and other members of the Molecular Cell Biomechanics Laboratory are gratefully acknowledged.

## Author contributions

N.D. and M.R.K.M. designed the study. N.D.. performed the simulations. G.Y., N.D., S.T. analyzed and interpreted the results. N.D., G.Y., S.T. drafted the manuscript. N.D. and M.R.K.M. revised the manuscript.

## Data & code availability

All data supporting this study are available as follows. Docking inputs/poses, MD input files, parameter/topology files, analysis scripts, and representative trajectories (compressed) will be deposited at Zenodo (https://doi.org/10.5281/zenodo.17957743).

---

**Figure S1.**
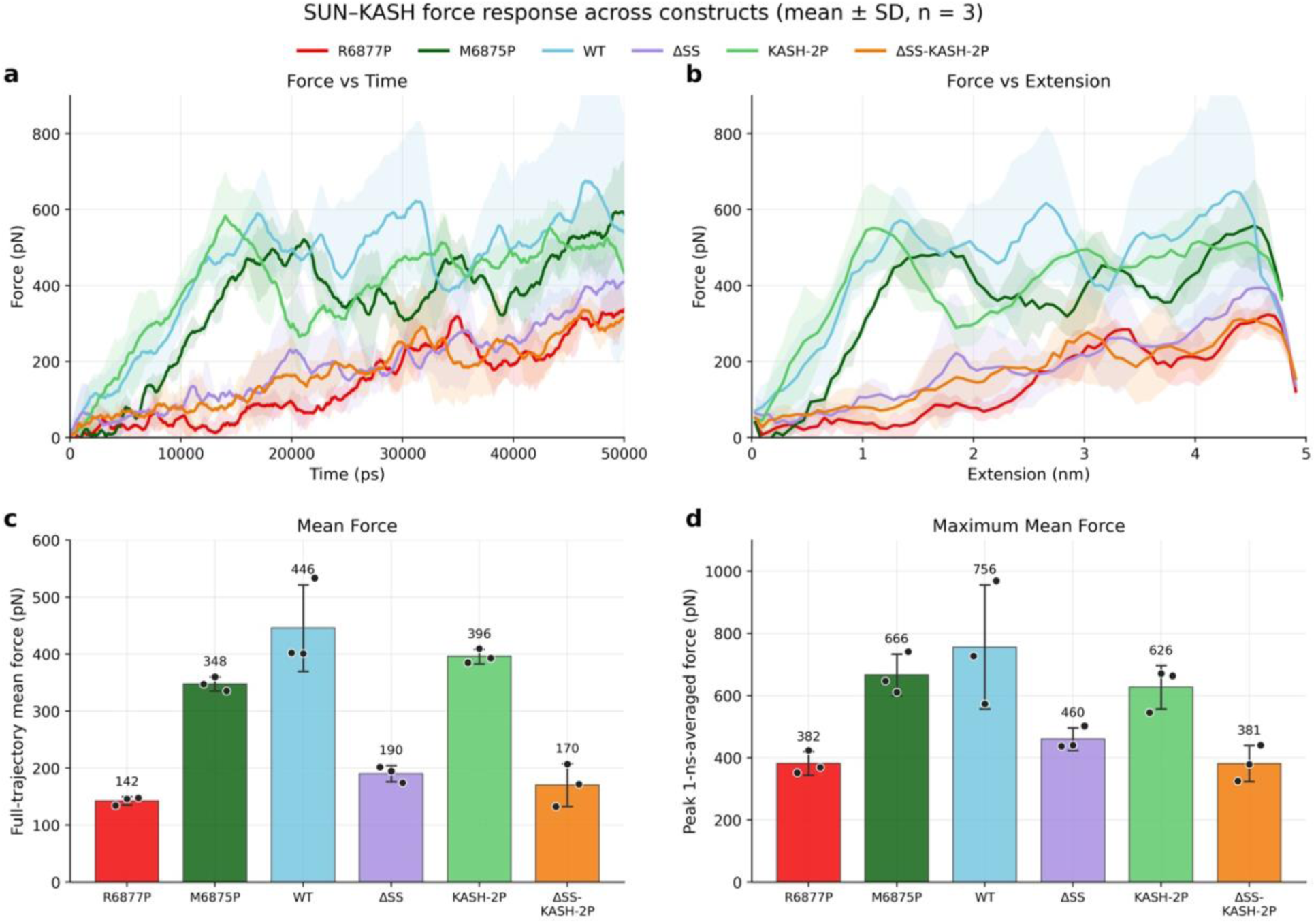
Force–time, force–extension, mean-force, and peak-force responses of single-point proline mutants compared with reference constructs. Two single-point proline substitutions were introduced to probe the mechanical roles of specific residues: ΔSS R6877P (red), in which arginine 6877 was replaced by proline in the disulfide-deleted (ΔSS) background, and M6875P (dark green), in which methionine 6875 was replaced by proline in the wild-type background. Reference constructs—WT (cyan), ΔSS (purple), KASH-2P (light green), and ΔSS-KASH-2P (orange)—are shown for comparison. (a) Force as a function of time. Force traces were smoothed using a centered 1-ns moving average. (b) Force as a function of extension. Force was binned by extension within each replicate before replicate averaging. In (a) and (b), solid lines represent the mean of three replicates and shaded regions represent ±1 SD. (c) Full-trajectory mean force. Each point represents the mean force from one 50-ns replicate; bars and error bars show the mean ± SD across three replicates. (d) Peak force, defined for each replicate as the maximum of its 1-ns-averaged force trace. Points represent replicate peak forces, and bars and error bars show the mean ± SD across three replicates. ΔSS R6877P transmits lower forces than ΔSS-KASH-2P during the early loading regime and subsequently converges toward similar force values, with overlapping replicate variability bands. M6875P shows delayed force onset relative to KASH-2P but reaches comparable forces at larger extensions. The bar-plot summaries similarly show that ΔSS R6877P has a lower full-trajectory mean force but a comparable peak force to ΔSS-KASH-2P, whereas M6875P has a lower mean force but a comparable peak force to KASH-2P. Neither single substitution produces the intermediate mechanical response expected if partial disruption of the β-sheet network resulted in proportionally greater force transmission than complete disruption.

